# Hypomethylation of *STAT1* and *HLA-DRB1* is associated with type-I interferon-dependent *HLA-DRB1* expression in lupus CD8+ T cells

**DOI:** 10.1101/398081

**Authors:** Shaylynn Miller, Patrick Coit, Elizabeth Gensterblum-Miller, Paul Renauer, Nathan C Kilian, Mark Schonfeld, Pei-Suen Tsou, Amr H Sawalha

**Affiliations:** Division of Rheumatology, Department of Internal Medicine, University of Michigan, Ann Arbor, MI, USA; Center for Computational Medicine and Bioinformatics, University of Michigan, Ann Arbor, MI, USA

**Keywords:** CD8+ T cells, lupus, epigenteic priming, HLA-DRB1, interferon

## Abstract

**Objective:** We examined genome-wide DNA methylation changes in CD8+ T cells from lupus patients and controls, and investigated the functional relevance of some of these changes in lupus.

**Methods:** Genome-wide DNA methylation of lupus and age, sex, and ethnicity-matched control CD8+ T cells was measured using the Infinium MethylationEPIC arrays. Measurement of relevant cell subsets was performed via flow cytometry. Gene expression was quantified by qPCR.

**Results:** Lupus CD8+ T cells had 188 hypomethylated CpG sites compared to healthy matched controls. Among the most hypomethylated were sites associated with *HLA-DRB1*. Genes involved in the type-I interferon response, including *STAT1*, were also found to be hypomethylated. IFNα upregulated HLA-DRB1 expression on lupus but not control CD8+ T cells. Lupus and control CD8+ T cells significantly increased STAT1 mRNA levels after treatment with IFNα. The expression of CIITA, a key interferon/STAT1 dependent MHC-class II regulator, is induced by IFNα in lupus CD8+ T cells, but not healthy controls. Co-incubation of naïve CD4+ T cells with IFNα-treated CD8+ T cells led to CD4+ T cell activation, determined by increased expression of CD69, in lupus patients but not in healthy controls. This can be blocked by neutralizing antibodies targeting HLA-DR.

**Conclusions:** Lupus CD8+ T cells are epigenetically primed to respond to type-I interferon. We describe an HLA-DRB1+ CD8+ T cell subset that can be induced by IFNα in lupus patients. A possible pathogenic role for CD8+ T cells in lupus that is dependent upon a high type-I interferon environment and epigenetic priming warrants further characterization.

## Introduction

Systemic lupus erythematosus (SLE) is a chronic relapsing autoimmune disease characterized by the production of autoantibodies and multiple organ involvement. The etiology of lupus is incompletely understood, however, heightened interest in changes specific to the DNA methylome of lupus lymphocytes is emerging [1-4]. Previous work examining differential DNA methylation in the T lymphocytes of lupus patients has primarily been performed with CD4+ T cells [5]. As lymphocytes are heavily involved in both the regulation and initiation of the immune response, investigation of DNA methylation changes in additional immunological cell types is of potential use to further elucidate unknown components of lupus pathogenesis.

While the epigenetic components of CD8+ T cells in lupus have yet to be described, the regulatory and functional changes of CD8+ T cells in lupus have been previously examined. Among total CD8+ T cells, lupus patients with active disease have increased proportion of naïve CD8+ T cells and reduced proportion of effector CD8+ T cells [6, 7]. Effector CD8+ T cells in lupus patients have reduced effector functionality through altered cytokine production, diminished suppressor function, and decreased cytotoxic T cell activity [6, 8]. The cytokine profiles of lupus CD8+ T cells have been found to favor increased IL-12 and decreased IL-6, thus resulting in dysregulation of the stimulatory and inhibitory roles of CD8+ T cells, respectively [8]. In addition, CD8+ T cells in lupus patients are characterized by reduced expression of signaling lymphocytic activation molecule family member 7 (SLAMF7), which is a type I transmembrane glycoprotein receptor that promotes effector CD8+ T cell function [6]. In contrast, Blanco et al. reported that patients with SLEDAI scores of seven or greater had a diminished naïve CD8+ T cell population and an elevated effector CD8+ T cell population [9]. CD8+ T cells are critical in blocking viral infections, which might trigger disease activation in lupus by increased type-I interferon production. Indeed, a pathogenic role for Epstein Barr virus infection in inducing lupus has been suggested, and linked to increased type-I interferon production, and more recently to genetic susceptibility in lupus [10-14]. As the role of CD8+ T cells in lupus remains incompletely understood, and is likely dependent upon currently unknown mechanisms, further examination of CD8+ T cell epigenetic changes in lupus could provide beneficial insight into this enigmatic disease.

In this study, we investigated genome-wide DNA methylation changes in the CD8+ T cells of lupus patients compared to age, sex, and ethnicity matched healthy controls. Functional annotation analysis of genes hypomethylated in lupus CD8+ T cells, followed by functional studies, suggest that lupus CD8+ T cells are epigenetically primed to respond to interferon (IFN) α and overexpress HLA-DRB1.

## Methods

### Patients and Controls

A total of 44 lupus patients (mean ± SEM age: 41.3 ± 1.6; median age: 42; and age range: 20 to 66 years) and 38 healthy controls (mean ± SEM age: 42.5 ± 1.9; median age: 40; and age range: 23 to 65 years) participated in this study. All lupus patients fulfilled the American College of Rheumatology classification criteria for SLE [15]. The mean SLEDAI score for lupus patients involved in this study was 3.57 with a median of 4 (range: 0 to 12). Lupus patients on cyclophosphamide or methotrexate were excluded from participating in the study as these drugs cause changes in cell surface expression of activation markers in lymphocyte subsets and altered epigenetic patterns, respectively [16-18]. All participants signed informed consent approved by the institutional review board of the University of Michigan.

### Sample Collection and DNA Extraction from Isolated CD8+ T cells

For DNA methylation studies, 8 lupus patients and 8 healthy controls recruited from the University of Michigan rheumatology clinics were matched by age (± 5 years), sex, and ethnicity (**Supplementary Table 1**). Peripheral blood mononuclear cells (PBMCs) were extracted from whole blood via Ficoll-Paque PLUS density centrifugation (GE Healthcare Bio-Sciences). Total T cells were isolated using negative selection by magnetic bead separation with the Human Pan T cell Isolation II kit (Miltenyi Biotec). Flow cytometry sorting was used to isolate CD3+CD56-TCRαβ+CD8+ T cells using the following fluorophore-conjugated antibodies: APC anti-human CD56 (clone: HCD56), FITC anti-human CD8a (clone: RPA-T8), Pacific Blue anti-human CD3 (clone: HIT3a), and PE anti-human alpha/beta-TCR (clone: IP26) (BioLegend), as previously described [19]. DNA was extracted from CD8+ T cells using the DNeasy Blood and Tissue kit (Qiagen) and bisulfite converted with the EZ DNA Methylation kit (Zymo Research) for DNA methylation analysis. It is worth noting that TCRαβ+ CD8+ T cells constituted ∼99% of CD8+ T cells in the peripheral blood of lupus patients or controls.

### Differential DNA Methylation and Gene Functional Annotation Analysis of CD8+ T cells in Lupus Patients

Methylation status of CD8+ T cells was measured by hybridizing bisulfite-converted DNA to the Infinium MethylationEPIC BeadChip array (Illumina). Methylation β values were calculated from raw intensity values using GenomeStudio Methylation module software (Illumina) after subtracting background, normalization to control probes, error correction using the Illumina custom model, and correction for false discovery rate. Group comparison and statistical significance were determined using Δ*β* = *β*_*SLE*_ *– β*_*control*_ and *DiffScore* = 10 × *sgn* (*β*_*SLE*_ – *β*_*control*_) × log_10_(*P*), respectively. Probes were filtered using the following criteria: |Δβ| ≥ 0.10, |DiffScore| ≥ 22 (equivalent to an FDR-adjusted *P* ≤ 0.01), detection *P*-value < 0.05, and no SNP within 10bp of the 3’ end of the probe. Gene annotations for hyper- and hypomethylated probes were used as input for gene function annotation analysis using DAVID [20]. Categories were restricted to molecular function and biological process gene ontologies and KEGG pathways, and only annotations with a minimum gene number of 2, a modified Fisher Exact P-value <0.1, and FDR ≤5% were considered.

### IFN-α treatment and Expression of HLA-DRB1 on CD8+ T cells

PBMCs were stimulated overnight in cell culture wells that were pre-coated with 10μg/ml of anti-human CD3 (Clone: UCHT1) and 2.5μg/ml of soluble anti-human CD28 (Clone: 28.2) (BD Biosciences) in T cell media (RPMI media with 10% fetal bovine serum and 2mM L-glutamine) in the presence of 1,000 U/ml IFNα 2B (Sigma-Aldrich). Culture media were then replaced and these cells were cultured for an additional four days with IFNα as this condition has been shown to produce peak HLA-DR expression on T cells [21]. At the end of the incubation, PBMCs were stained with fluorochrome-conjugated antibodies for the cell-surface markers CD3, CD8a, and HLA-DRB1. The detailed flow cytometry procedure and analysis is described below.

### Co-incubation of IFN-α Treated CD8+ T cells with Naïve CD4+ T cells

After extracting from whole blood through Ficoll-Paque PLUS density centrifugation (GE Healthcare Bio-Sciences), PBMCs were divided into three portions. The first portion was cultured in T cell media for 5 days without any stimulation and treatment for naïve CD4+T cell isolation using the Naïve CD4+ T Cell Isolation Kit II (Miltenyi Biotec), which allows for isolating untouched naïve CD4+ T cells. The second portion was stimulated with anti-human CD3 and anti-human CD28 in the presence of 1,000 U/ml IFN-α 2B (Sigma-Aldrich) overnight in T cell media. These cells were then cultured for an additional four days with IFNα. The last portion of the PBMCs were similarly stimulated overnight with anti-human CD3 and anti-human CD28, and cultured for a total of 5 days, however, without IFNα treatment. At the end of the culture period, CD8+ T cells were isolated from these two portions via the CD8+ T Cell Isolation Kit (Miltenyi Biotec), which isolates untouched TCRαβ+ CD8+ T cells. To examine whether IFNα-treated CD8+ T cells can stimulate naïve CD4+ T cells, the isolated CD8+ T cells with or without IFNα treatment were incubated with naïve CD4+ T cells for 20-22 hours in a 1:1 ratio. The cells were then stained with fluorochrome-conjugated antibodies for CD3, CD4, and CD69 at a concentration of 2 × 10^4^ cells per 1uL of 1X PBS. The percentage of CD4+ T cells bearing CD69, an early activation marker, was measured by flow cytometry for the CD3+ CD4+ T cells of both co-incubation groups, as described below.

### Co-incubation of IFNα treated CD8+ T cells with Naïve CD4+ T cells in the Presence of anti-HLA-DR Blocking Antibody

PBMCs were isolated and divided into two groups. The first group was incubated in T cell media for five days and naïve CD4+ T cells were isolated. PBMCs in the second group were activated overnight with anti-human CD3 and anti-human CD28, and treated with 1,000 U/ml IFNα overnight. The media for the PBMCs were then replaced and the cells were treated with IFNα for an additional four days. At the end of the incubation period, untouched TCRαβ+ CD8+ T cells were isolated using the CD8+ T Cell Isolation Kit (Miltenyi Biotec) and split evenly into three groups: Anti-HLA-DR, isotype control, and no antibody. Prior to use, the Fc portions of the LEAF™ Purified anti-human HLA-DR Antibody and LEAF™ Purified Mouse IgG2a, κ Isotype Ctrl Antibody were degraded using the Pierce™ F(ab’)2 Preparation Kit (Thermo Fisher Scientific) to diminish non-specific binding of the Fc portion. The CD8+ T cells were treated with either 20ug/ml of LEAF™ Purified anti-human HLA-DR Antibody (Clone: L243) or 20ug/ml of LEAF™ Purified Mouse IgG2a, κ Isotype Ctrl Antibody (Clone: MOPC-173) (BioLegend,) at 37°C for 1 hour, while the CD8+ T cells in the no antibody group were left untreated. The cells were then washed with PBS twice to remove any residual IgG. The CD8+ T cells from the three groups were then co-incubated with naïve CD4+ T cells overnight. At the end, cells were stained with fluorochrome-conjugated antibodies for CD3, CD4, and CD69. The percentage of CD4+ T cells expressing CD69, an early activation marker, was measured by flow cytometry for the CD3+ CD4+ T cells for all co-incubation groups.

### Flow Cytometry Analysis

PBMCs were stained with fluorochrome-conjugated antibodies at a concentration of 2×10^4^ cells per 1uL of 1X PBS for 30 minutes on ice. The following fluorochrome-conjugated antibodies were used for flow cytometry antibody staining: FITC anti-human CD8a (clone: RPA-T8), APC anti-human CD69 (clone: FN50), APC/Cy7 anti-human CD4 (clone: RPA-T4), Pacific Blue anti-human CD3 (clone: UCHT1), and PE anti-human HLA-DRB1 (clone: NFLD.D2). All fluorochrome-conjugated antibodies were purchased from BioLegend. PBMCs were fixed with 500uL of Biolegend Fixation Buffer per 10^6^ cells for 20 minutes at ambient temperature. The iCyt Synergy SY3200 Cell Sorter (Sony Biotechnology Inc.) was used in conjunction with WinList 9.0.1 software (Verity Software House) for flow cytometry sample processing and analysis, respectively. PBMCs were gated to select for CD3+ CD8+ HLA-DBR1+ T cells. The median fluorescence intensity (MFI) of PE anti-human HLA-DRB1 was measured for the CD3+ CD8+ T cell population and normalized to the background PE anti-human HLA-DRB1 MFI of the unstained PBMC population. The normalized MFI (nMFI) was determined by the following equation 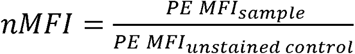 [22]. FlowJo version 10 was used to generate MFI histograms (FlowJo LLC).

### Quantitative Real-time PCR

RNA was isolated with the Direct-zol RNA MiniPrep kit according to the manufacturer’s instructions (Zymo Researcg). The Verso cDNA Synthesis Kit (Thermo Fisher Scientific) was used along with reverse transcriptase PCR to produce the cDNA necessary for quantitative real-time PCR (qPCR). qPCR was done with Power SYBR™ Green PCR Master Mix (Thermo Fisher Scientific) and used along with human KiCqStart SYBR Green primers to measure mRNA expression of *STAT1* and *CIITA* (Sigma-Aldrich). All gene targets were run in duplicates and normalized to *β-actin*. The ABI 7900HT instrument along with SDS v2.4 software was used for qPCR.

### Statistical Analysis

Graphpad Prism 7.0 was used for the purposes of statistical analysis and creation of graphs. For comparing sample groups from the same subjects but with different cell culture treatments the paired, non-parametric Wilcoxon Test was used. For tests comparing the same treatment conditions but with different subjects the unpaired, non-p arametric, Mann-Whitney Test was used. For all statistical tests, a significance cutoff of *P* < 0.05 was used.

## Results

### DNA methylation changes in lupus CD8+ T cells

DNA methylation analysis of lupus CD8+ T cells revealed significant hypermethylation of 943 loci and hypomethylation of 188 loci compared to matched control CD8+ T cells (**Supplementary Table 2**). The most hypomethylated CpG site was found within the major histocompatibility complex class II gene *HLA-DRB1* (cg11404906; Δβ=-0.333) (**Table 1**). Four other loci were hypomethylated in *HLA-DRB1*, two of which were also among the most hypomethylated (cg09949906; Δβ= −0.270 & cg09139047; Δβ=-0.247). These loci are in or around exon 2 of *HLA-DRB1* and a DNase I hypersensitivity region that suggest a regulatory region that can be open and available to transcription factors and machinery (**Figure 1**). The most hypermethylated site was not associated with a gene (cg01005486; Δβ=0.325), but loci associated with *HLA-DRB5* (cg22635523; Δβ=0.239) and *HLA-DRB6* (cg06559318; Δβ=0.207 & cg20083297; Δβ=0.189) did have significantly higher methylation levels in lupus CD8+ T cells (**Table 1**).

**Table 1:** Top 25 hypomethylated and hypermethylated loci in lupus CD8+ T cells.

| Probe ID | $\Delta\beta$ | Patient Average Methylation ($\beta$) | Control Average Methylation ($\beta$) | Genomic Locus (chr:position) (hg19) | DiffScore | Gene |
| --- | --- | --- | --- | --- | --- | --- |
| cg11404906 | -0.333 | 0.302 | 0.635 | 6:32551749 | -337.887 | *HLA-DRB1* |
| cg02453376 | -0.330 | 0.402 | 0.732 | 1:152586240 | -337.887 | *LCE3B* |
| cg14392283 | -0.319 | 0.445 | 0.765 | 8:144103587 | -337.887 | *LY6E* |
| cg10978274 | -0.317 | 0.412 | 0.729 | 1:152572665 | -337.887 | *LCE3C* |
| cg10829391 | -0.308 | 0.418 | 0.725 | 14:101069717 | -337.887 |  |
| cg09949906 | -0.270 | 0.361 | 0.630 | 6:32552350 | -250.225 | *HLA-DRB1* |
| cg00293330 | -0.266 | 0.340 | 0.606 | 1:152572678 | -242.598 | *LCE3C* |
| cg17748470 | -0.252 | 0.421 | 0.674 | 11:4969161 | -219.081 | *OR51A4* |
| cg12393503 | -0.252 | 0.473 | 0.725 | 3:172468831 | -232.588 |  |
| cg09139047 | -0.247 | 0.462 | 0.709 | 6:32552042 | -217.006 | *HLA-DRB1* |
| cg16180556 | -0.245 | 0.286 | 0.531 | 1:110230269 | -204.564 | *GSTM1* |
| cg21549285 | -0.239 | 0.469 | 0.707 | 21:42799141 | -201.317 | *MX1* |
| cg26184673 | -0.221 | 0.568 | 0.789 | 12:131520095 | -201.400 | *ADGRD1* |
| cg08822897 | -0.215 | 0.348 | 0.564 | 11:64258103 | -144.881 |  |
| cg27244972 | -0.212 | 0.418 | 0.630 | 21:47716529 | -140.917 | *C21orf57* |
| cg02484732 | -0.207 | 0.361 | 0.569 | 6:56096198 | -132.102 | *COL21A1* |
| cg20684491 | -0.204 | 0.555 | 0.758 | 1:25596433 | -156.792 |  |
| cg09234453 | -0.199 | 0.375 | 0.575 | 6:18402672 | -120.179 | *RNF144B* |
| cg21158163 | -0.198 | 0.681 | 0.879 | 20:59542578 | -229.793 |  |
| cg16361921 | -0.198 | 0.384 | 0.582 | 5:106599492 | -118.102 |  |
| cg06650861 | -0.197 | 0.379 | 0.576 | 4:169238926 | -116.972 | *DDX60* |
| cg01609658 | -0.196 | 0.200 | 0.396 | 1:151694478 | -143.422 | *RIIAD1* |
| cg07815522 | -0.194 | 0.480 | 0.675 | 3:122282157 | -120.319 | *PARP9;DTX3L* |
| cg24311634 | -0.192 | 0.557 | 0.749 | 12:131519883 | -134.197 | *ADGRD1* |
| cg18220841 | -0.188 | 0.344 | 0.531 | 16:88160246 | -104.890 |  |
| cg11102724 | 0.182 | 0.293 | 0.112 | 5:150619039 | 340.551 |  |
| cg00256329 | 0.186 | 0.702 | 0.515 | 17:724374 | 114.220 | *NXN* |
| cg24432675 | 0.186 | 0.603 | 0.417 | 10:1505472 | 102.562 | *ADARB2* |
| cg19214707 | 0.187 | 0.406 | 0.219 | 7:3157722 | 125.122 |  |
| cg27246321 | 0.189 | 0.462 | 0.273 | 2:165924708 | 114.220 |  |
| cg02419513 | 0.189 | 0.299 | 0.110 | 8:90255978 | 340.551 |  |
| cg20083297 | 0.189 | 0.685 | 0.496 | 6:32528390 | 114.865 | *HLA-DRB6* |
| cg09398595 | 0.190 | 0.751 | 0.561 | 14:98255618 | 340.551 |  |
| cg24919907 | 0.193 | 0.799 | 0.606 | 14:98255601 | 340.551 |  |
| cg14507403 | 0.196 | 0.550 | 0.355 | 7:97798005 | 115.218 | *LMTK2* |
| cg11244641 | 0.207 | 0.591 | 0.384 | 2:241343025 | 340.551 |  |
| cg06559318 | 0.207 | 0.728 | 0.521 | 6:32526260 | 340.551 | *HLA-DRB6* |
| cg10123377 | 0.212 | 0.584 | 0.373 | 3:42387524 | 340.551 |  |
| cg12022450 | 0.213 | 0.687 | 0.474 | 22:24353218 | 340.551 |  |
| cg21787089 | 0.213 | 0.806 | 0.593 | 3:46484250 | 340.551 | *LTF* |
| cg01427108 | 0.225 | 0.668 | 0.443 | 3:46484379 | 340.551 | *LTF* |
| cg25755428 | 0.227 | 0.570 | 0.343 | 19:13875111 | 340.551 | *MRI1* |
| cg14188831 | 0.233 | 0.299 | 0.066 | 1:25594523 | 340.551 |  |
| cg09232555 | 0.236 | 0.508 | 0.272 | 8:11619866 | 340.551 |  |
| cg22635523 | 0.239 | 0.682 | 0.443 | 6:32492390 | 340.551 | *HLA-DRB5* |
| cg05176970 | 0.252 | 0.707 | 0.455 | 17:724273 | 340.551 | *NXN* |
| cg12108278 | 0.291 | 0.892 | 0.601 | 17:15789774 | 340.551 |  |
| cg12340462 | 0.297 | 0.650 | 0.353 | 9:46141878 | 340.551 |  |
| cg18213661 | 0.321 | 0.722 | 0.401 | 11:93681423 | 340.551 |  |
| cg01005486 | 0.325 | 0.802 | 0.477 | 3:13246006 | 340.551 |  |

**Figure 1:**
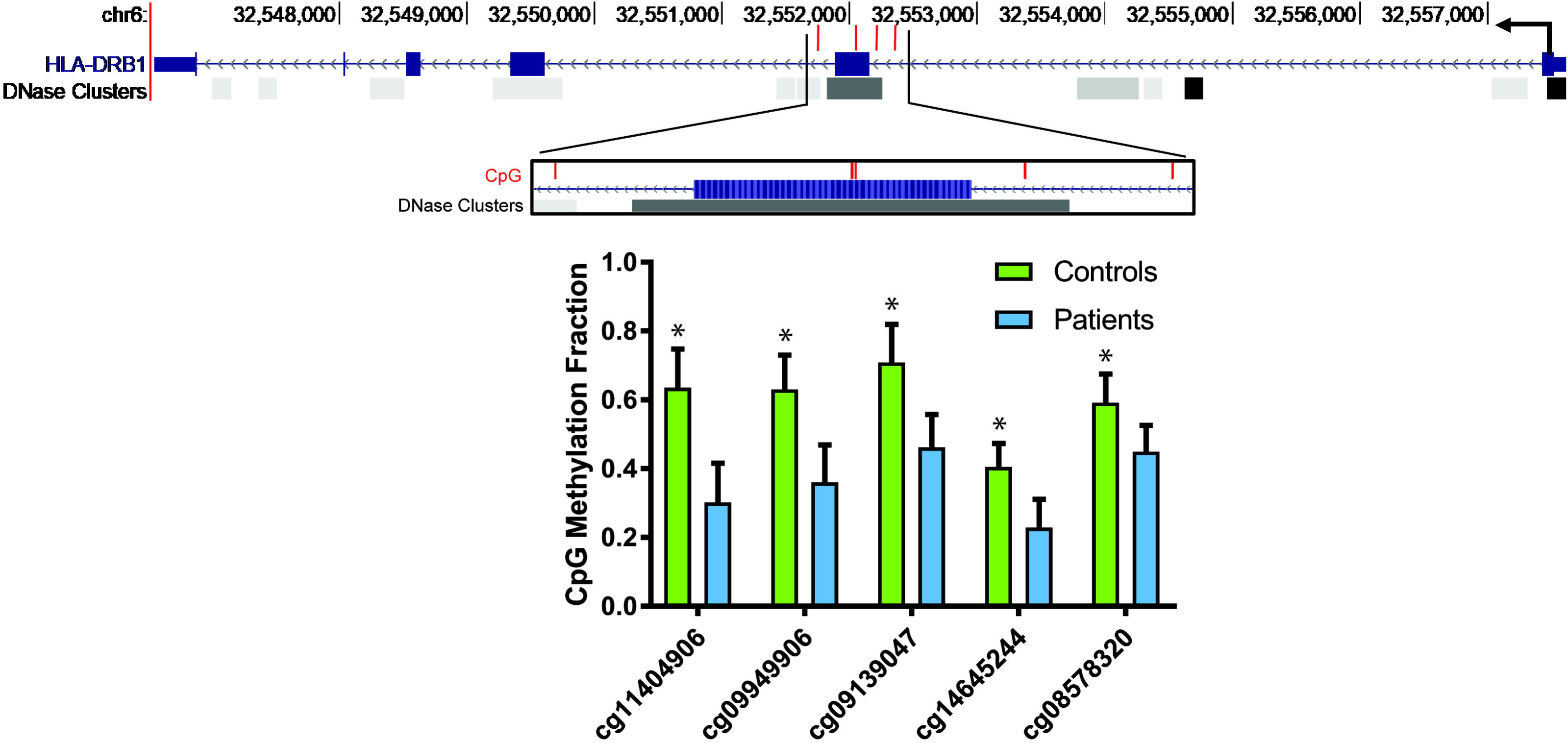
Diagram depicting the *HLA-DRB1* gene and the positions of the 5 CpG sites within *HLA-DRB1* hypomethylated in lupus CD8+ T cells compared to normal healthy controls (each CpG site is shown with a red bar). The hypomethylated region overlaps with a known DNase hypersensitivity cluster, suggesting a regulatory potential. “HLA-DRB1” and “DNase Clusters” tracks were retrieved from UCSC Genome Browser. DNase Clusters represent regions of DNase I hypersensitivity as described in the “DNaseI Hypersensitivity Clusters in 125 cell types from ENCODE (V3)” track [33]. Lower panel shows DNA methylation fractions on each of the CpG sites in patients and controls (mean±SEM). *P<0.01 after correction for multiple testing for all probes on the array.

Genes involved in the type I interferon response were also found to be hypomethylated in lupus CD8+ T cells including the transcription factor *STAT1* (cg00676801; Δβ=-0.150 & cg14951497; Δβ=-0.138) and *MX1* (cg21549285; Δβ=-0.239 & cg22862003; Δβ=- 0.122) (**Table 1 and Supplemental Table 2**), both type I interferon-related genes that have previously been identified as being hypomethylated in several immune cell types in lupus patients, including CD4+ T cells [1]. This was supported by gene enrichment analysis, which revealed a predominance of gene ontology terms for interferon response and signaling: “interferon-gamma-mediated signaling pathway” (GO:0060333; *P*=1.41E-05), “defense response to virus” (GO:0051607; *P*=7.56E-04) and “type I interferon signaling pathway” (GO:0060337; P=2.72E-03) (**Supplemental Table 3**). Antigen presentation was also a major theme that included “peptide antigen binding” (GO: 0042605; *P*=2.17E-04) and “antigen processing and presentation” (GO:0019882; *P*=1.76E-03) terms. KEGG pathways enriched in hypomethylated genes showed that “Cell adhesion molecules (CAMs)” (KEGG:hsa04514; *P*=6.73E-05) was the most significantly enriched pathway among hypomethylated genes (**Supplemental Table 3**). Several disease pathways were also enriched including autoimmune diseases like rheumatoid arthritis (KEGG:hsa05323; *P*=9.51E-04) and systemic lupus erythematosus (KEGG:hsa05322; *P*=4.44E-03) as well as viral infections (“Influenza A” KEGG:hsa05164; *P*=1.68E-03 (“Herpes simplex infection” KEGG:hsa05168; *P*=2.10E-03). A pathway including glutathione *S*-transferase genes (GST) (’Drug metabolism – cytochrome P450” KEGG:hsa00982; *P*=4.76E-03) was also enriched (**Supplemental Table 3**). GSTs catalyze the conjugation of reduced glutathione to electrophilic compounds like xenobiotics and increase their solubility and excretion from the cell and reduce the toxicity of reactive oxygen species in the cell [23]. The hypermethylated genes displayed broader functional annotation groups and did not show dominance of enriched pathways by HLA genes that were seen among hypomethylated genes (**Supplemental Table 3**).

### IFNα induces HLA-DRB1 expression on lupus CD8+ T cells

*HLA-DRB1* was robustly hypomethylated in lupus CD8+ T cells; we also observed hypomethylation in *STAT1* and the interferon-signaling pathway, which regulate MHC class II expression. Therefore, we sought to study the functional implications of these epigenetic changes by investigating the expression of HLA-DRB1 and the role of type-I interferon response in the CD8+ T cells of lupus patients. Indeed, we found that both patients and controls possess a CD8+HLA-DRB1+ T cell subset, however, the subset’s response to IFNα differs in lupus patients. In lupus CD8+ T cells treated with IFNα, the expression of HLA-DRB1 was significantly elevated compared to untreated cells (P=0.02). In addition, when lupus CD8+ T cells were treated with IFNα, they expressed significantly more HLA-DRB1 on their surface than IFNα treated healthy control cells (p=0.04, **Figure 2A and B**). Similarly, HLA-DRB1 expression, as measured by normalized median fluorescence intensity, was increased with IFNα treatment in lupus CD8+ T cells (P= 0.04), but not in healthy controls (P=0.79, **Figure 2C and D**). These data suggest that hypomethylation of *HLA-DRB1* epigenetically primes lupus CD8+ T cells for type-I interferon dependent HLA-DRB1 overexpression.

**Figure 2:**
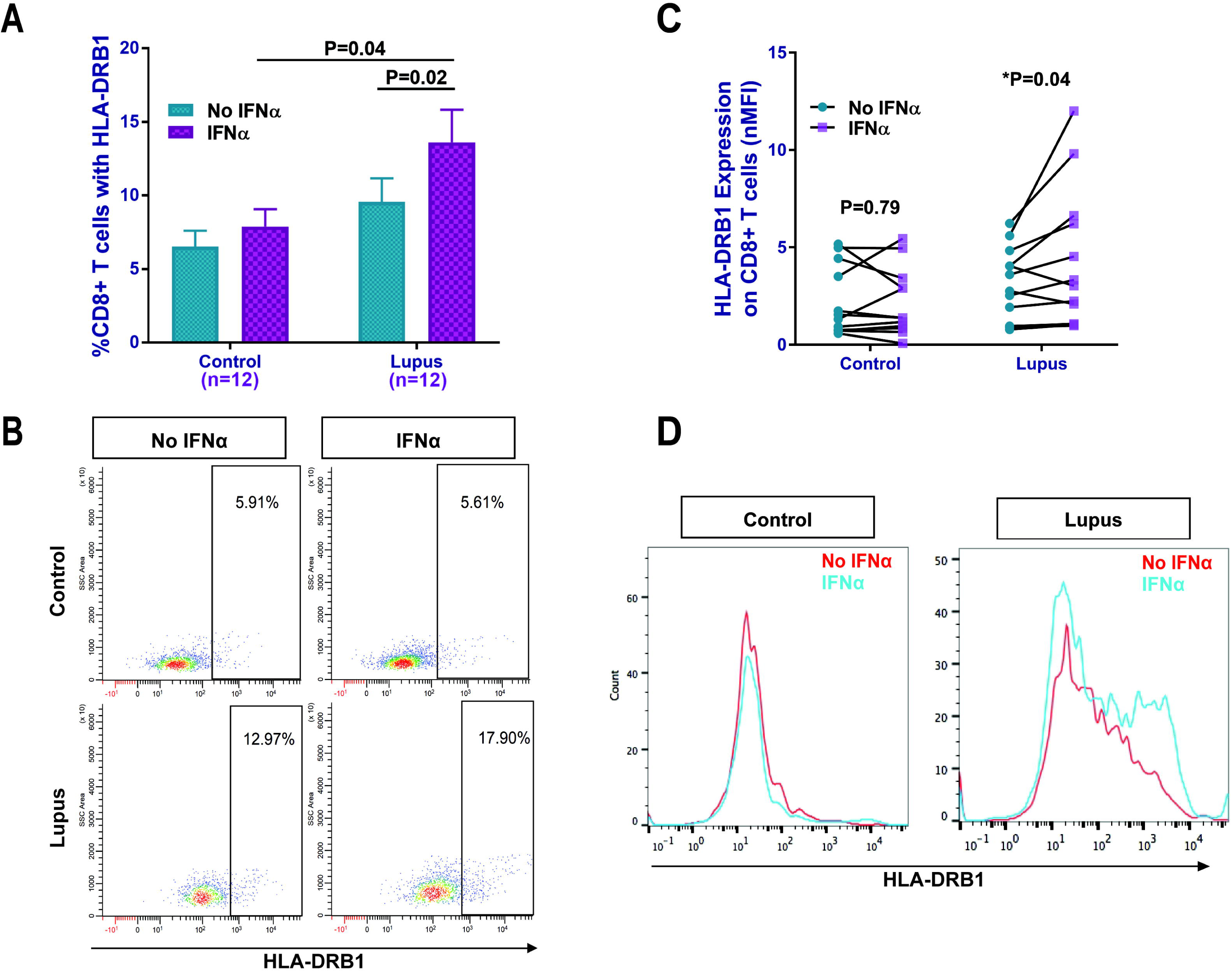
IFNα significantly increased HLA-DRB1 expression on lupus CD8+ T cells but not in healthy controls. **(A)** The percentage of CD8+ T cells expressing HLA-DRB1 in lupus patients and healthy controls with and without IFNα, and a representative dot plot **(B)**. Normalized median fluorescence intensity (nMFI) data **(C)**, and **(D)** representative histograms (MFI) from a lupus patient and a healthy control with and without IFNα, showing increased HLA-DRB1 expression with IFNα on the cell surface of lupus CD8+T cells. Results are presented as mean±SEM, and p<0.05 was considered significant. N=number of subjects.

### STAT1 and CIITA are elevated in IFNα-treated lupus CD8+ T cells

We observed hypomethylation of loci associated with STAT1 in lupus CD8+ T cells (**Supplemental Table 2**). STAT1 and CIITA are both important regulators in the MHC-II expression pathway (**Supplemental Figure 1**). For these reasons, we evaluated the expression of *CIITA* and *STAT1* mRNA in lupus CD8+ T cells with and without IFNα treatment. *STAT1* expression was elevated following IFNα treatment in both lupus patients (P= 0.02) and controls (P= 0.02) (**Figure 3A**). The expression of *CIITA* was also induced by IFNα stimulation in lupus CD8+ T cells, however, not in healthy controls (P=0.03) (**Figure 3B**). These findings suggest a type-I IFN dependent mechanism for the increased expression of HLA-DRB1 on lupus CD8+ T cells possibly mediated through *STAT1* and *CIITA* in a high type-I IFN environment.

**Figure 3:**
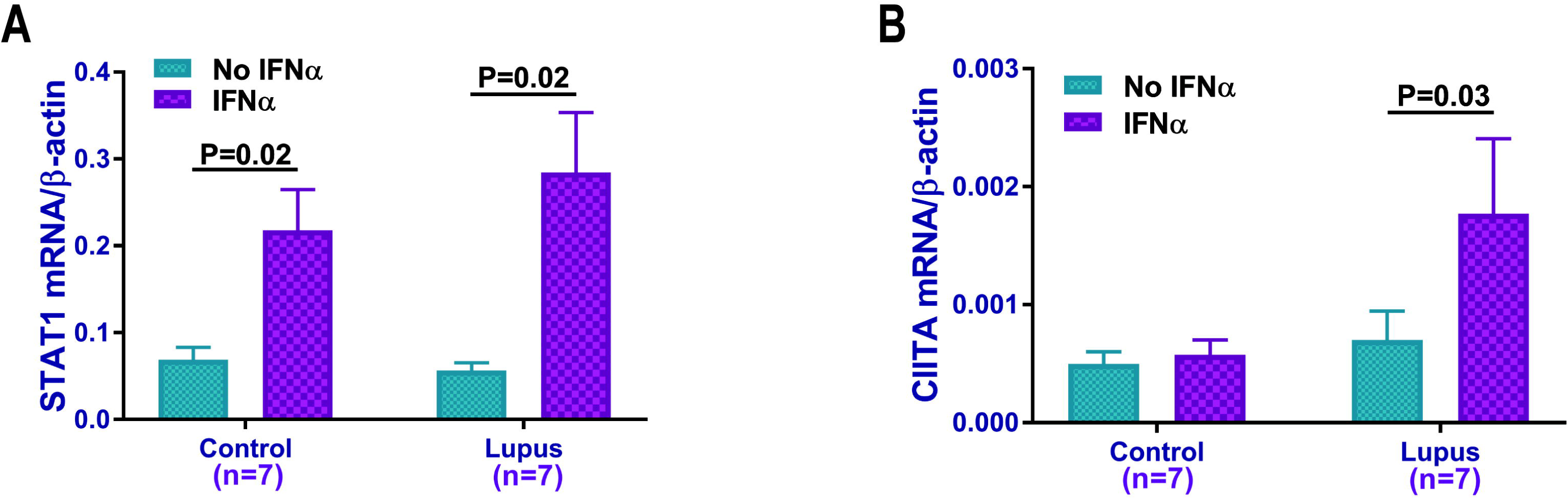
*STAT1* and *CIITA* mRNA expression in CD8+ T cells**. (A)** IFNα treatment significantly increased *STAT1* mRNA expression in CD8+ T cells from both lupus patients and healthy controls. **(B)** IFNα treatment significantly increased *CIITA* mRNA expression in CD8+ T cells from lupus patients but not healthy controls. Results are presented as mean±SEM, and p<0.05 was considered significant. N=number of subjects.

### Lupus CD8+ T cells activate autologous naïve CD4+ T cells in a type-I interferon and HLA-DR regulated manner

We showed that *HLA-DRB1* and *STAT1* are hypomethylated in lupus CD8+ T cells, which primes for increased HLA-DRB1 cell surface expression in the presence of IFNα. To demonstrate if HLA-DRB1 overexpression on lupus CD8+ T cells has any functional consequences, we examined whether lupus CD8+ T cells have the ability to stimulate autologous naïve CD4+ T cells. IFNα treatment increased the ability of lupus CD8+ T cells to induce CD69 expression, which is an early marker for T cell stimulation, on autologous naïve CD4+ T cells (p=0.002) while it had non-significant minimal effect on CD8+ T cells from healthy controls (**Figure 4A and B)**. This difference in response resulted in significant elevation of the percentage of stimulated CD4+ T cells after incubation with IFNα-treated CD8+ T cells from lupus patients compared to those isolated from healthy controls (P=0.04). Similarly, significant elevation of CD69 expression, as measured by nMFI, on CD4+ T cells after co-incubation with IFNα - treated lupus CD8+ T cells was also observed compared to IFNα-treated control CD8+ T cells (P=0.0004; **Supplemental Figure 2**).

**Figure 4:**
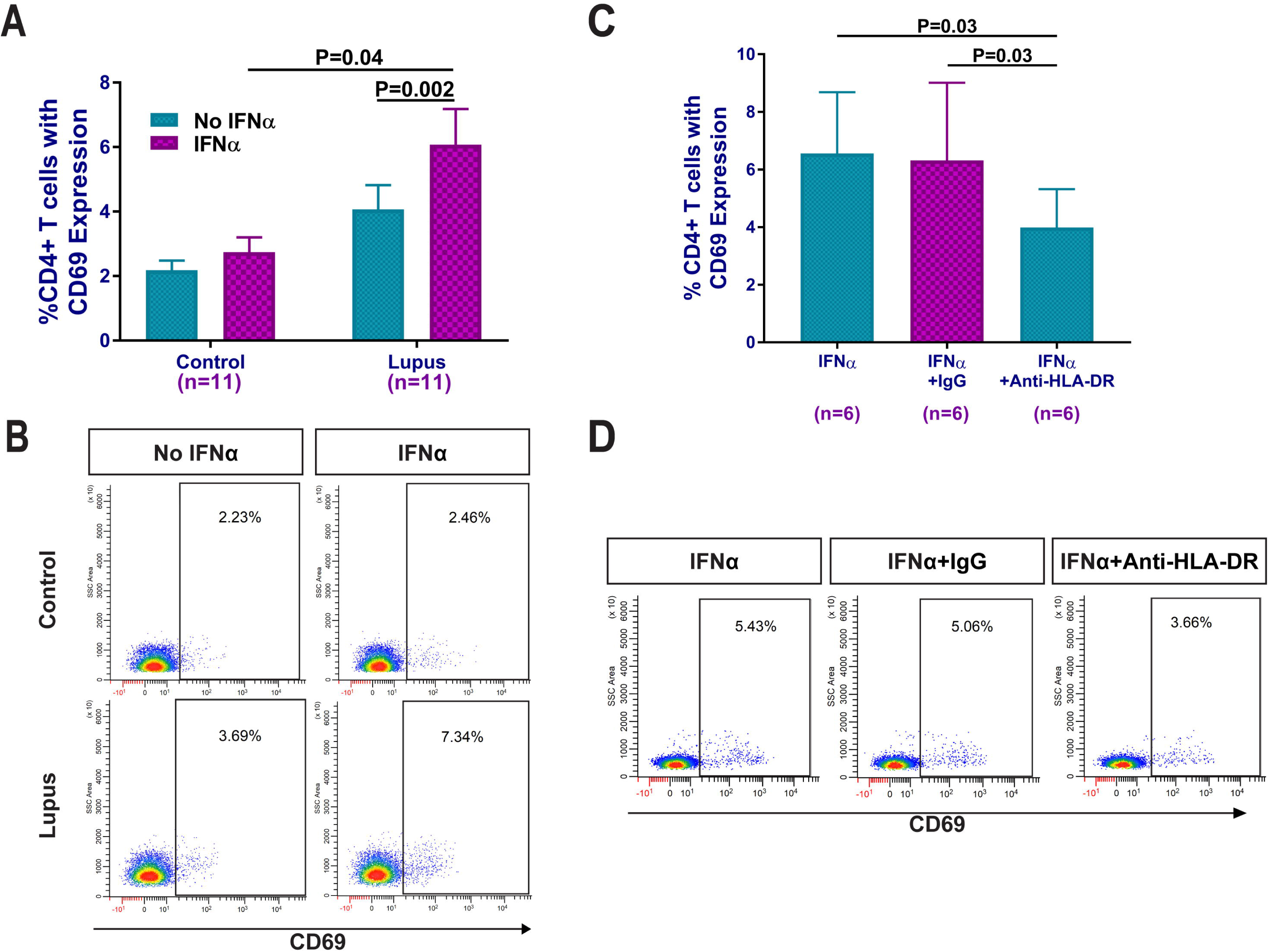
Effect of CD8+ T cells on activating autologous naïve CD4+ T cells. **(A)** Lupus and control CD8+ T cells, with and without IFNα, were co-cultured with autologous naïve CD4+ T cells and the percentage of CD3+CD4+CD69+ T cells were evaluated. A representative dot plot is shown in panel **(B)**. Blocking HLA-DR on IFNα-treated lupus CD8+ T cells abrogates co-stimulation of naïve CD4+ T cells compared to isotype IgG control **(C)**. A representative dot plot of 6 independent experiments is shown in **(D).** Results are presented as mean±SEM, and p<0.05 was considered significant. N=number of subjects.

IFNα treatment increases HLA-DRB1 expression on lupus CD8+ T cells. To determine if activation of naïve CD4+ T cells in the co-culture experiments is dependent upon HLA-DRB1 expression on IFNα-treated CD8+ T cells, we performed blocking experiments using neutralizing antibodies to HLA-DR. The increase in CD3+CD4+ CD69+ T cells as a result of co-incubation with IFNα-treated lupus CD8+ T cells was inhibited when CD8+T cells were pre-treated with anti-HLA-DR blocking antibody (**Figure 4C and D**, P=0.03). These data suggest that IFNα-treated lupus CD8+ T cells are capable of stimulating autologous naïve CD4+ T cells in a manner dependent upon induced HLA-DRB1 expression.

## Discussion

We performed a DNA methylation analysis in lupus CD8+ T cells and identified robust hypomethylation in CpG sites associated with *HLA-DRB1* and *STAT1* compared to healthy matched controls. Because Interferon/STAT1 signaling induces MHC class II expression, and type-I interferon production is increased in lupus patients, we hypothesized that demethylation in *HLA-DRB1* and *STAT1* might be associated with HLA-DRB1 expression on lupus CD8+ T cells, especially in the presence of type-I interferon. Indeed, our data demonstrate HLA-DRB1 cell surface expression in a subset of CD8+ T cells that is expanded in lupus compared to healthy controls, and that HLA-DRB1 expression can be induced in the presence of type-I interferon in lupus but not control CD8+ T cells. These data suggest that epigenetic priming facilitates a type-I interferon-dependent overexpression of HLA-DRB1 in lupus CD8+ T cells. We further show that lupus CD8+ T cells treated with type-I interferon can activate autologous naïve CD4+ T cells, and that this activation is abrogated by pre-incubating lupus CD8+T cells with HLA-DR blocking antibodies.

Both *HLA-DRB1* and *STAT1,* which are hypomethylated in lupus CD8+ T cells, are two possible down-stream gene targets of the JAK-STAT signaling cascade initiated by type-I IFN signaling (**Supplemental Figure 1**) [24-26]. IFNα can upregulate the surface expression of HLA-DRB1 by first binding to an IFNα receptor on the cell surface, and initiate the JAK-STAT signaling cascade that produces STAT1-STAT1 (STAT1-1) homodimers and STAT1-STAT2 (STAT1-2) heterodimers [27, 28]. STAT1-1 homodimers and STAT1-2 heterodimers translocate to the nucleus where they bind to the gamma-activated sequence (GAS) and allow for the expression of the class II transactivator (CIITA) transcription factor; CIITA binds to the MHC-II enhanceosome which upregulates MHC-II expression [24-26]. The canonical route taken for STAT dimerization during a type-I IFN response is a STAT1-2 heterodimer binding to ISGF3, translocating to the nucleus, and binding to the Interferon Stimulated Response Element (ISRE) to initiate the antiviral response [27]. However, as STAT1 homodimers binding to GAS are the typical path necessary for MHC-II expression, it is important to note that the type-I IFN response can also produce STAT1 homodimers and STAT1-2 heterodimers that can bind to GAS [24, 28, 29]. In this study we indeed demonstrated that type-I IFN stimulates *STAT1* expression in CD8+ T cells to a similar extent in both healthy controls and lupus patients. However, the response in *CIITA* expression after type-I IFN treatment is what sets the controls and patients apart; *CIITA* was significantly elevated after IFNα treatment in lupus CD8+ T cells but not in healthy controls. This might potentially explain the higher HLA-DRB1 expression on lupus CD8+ T cells.

We also identified a subset of cell adhesion molecules differentially methylated in lupus CD8+ T cells. *SELL* encodes the cell adhesion molecule L-selectin (Lymphocyte-selectin) and was hypomethylated at a single locus in our analysis (cg06816239; Δβ=-0.15). L-selectin is vital to the ability of activated CD8+ T cells to traffic to tissues infected by viruses [30]. Our analysis also revealed that a single CpG site in the 3’ untranslated region of *SELPLG* was hypermethylated in lupus CD8+ T cells (cg02520593; Δβ= 0.17). PSGL-1 (encoded by *SELPLG*) is a ligand for the P-selectin that plays a role in T cell tissue infiltration [31]. PSGL-1 expression is associated with reduced TCR and IL-2 stimulatory activity in murine CD8+ T cells, leading to decreased cell survival and tumor cell killing [31]. Despite the enrichment in cell adhesion molecules that was observed from the DNA methylation profiles, we did not observe any significant differences in cell adhesion between control and lupus CD8+ T cells in a T cell-endothelial cell assay (data not shown).

Previous studies investigating the role of CD8+ T cells in lupus have primarily focused on T cell cytotoxicity, autoantibody production, and cellular markers indicative of disease flares. Most reports showed defective cytotoxic responses in lupus CD8+ T cells, which has been suggested as a possible mechanism contributing to increased infection rates in lupus patients [6, 8]. Other studies, however, suggest that patients with higher disease activity may have increased cytotoxic T cell activation and elevated proportion of CD8+ T cells expressing granzyme B, perforin, and HLA-DR [9, 32]. In our study, we did not detect DNA methylation changes in the gene loci encoding for perforin and granzyme B. In addition, other cytotoxic gene loci encoding for granzyme A, granzyme H, and granzyme K, granzyme M, and the genes encoding transcription factors eomesodermin (eomes) and T-bet, which play key roles in CD8+ T cell differentiation, were also not differentially methylated between lupus patients and controls.

In summary, we describe DNA methylation changes in lupus CD8+ T cells at a genome-wide level, and provide evidence that CD8+ T cells in lupus are epigenetically primed for HLA-DRB1 overexpression in response to type-I IFN. A CD8+ HLA-DRB1+ T cell subset is expanded in lupus patients compared to healthy controls, and can be induced by IFNα in lupus but not healthy control CD8+ T cells. In lupus patients, CD8+HLA-DRB1+ T cells can stimulate autologous naïve CD4+ T cells in a manner dependent upon HLA-DRB1 expression. Our data link a higher type-I interferon environment in lupus patients to a pathogenic CD8+ T cell subset, which can be potentially abrogated by blocking type-I interferon response. These findings provide novel insights into possible pathogenic roles for CD8+ T cells in lupus, and suggest an interaction between high type-I interferon environment in lupus patients with epigenetic dysregulation in CD8+ T cells. Additional studies to evaluate if lupus-associated genetic variants within the HLA region influence DNA methylation patterns in the *HLA-DRB1* locus are warranted.

## Acknowledgement

This work was supported by the National Institute of Allergy and Infectious Diseases of the National Institutes of Health grant number R01AI097134. We would like to thank Ye He for her help with flow cytometry analysis.

## Notes

**Conflict of interest:** None of the authors has any financial conflict of interest to disclose

